# Fluffy nests and bloody parasites: feather lining does not mitigate bird blow flies infestations

**DOI:** 10.64898/2026.10.04.756503

**Authors:** Simon Coroller-Chouraki, Jade Savage, Fanie Pelletier, Dany Garant, Marc Bélisle

## Abstract

Feathers incorporated into bird nests are thought to provide multiple benefits, including thermal insulation and protection against nest-dwelling ectoparasites. For instance, in Swallows, it has been suggested that feathers influence infestations by *Protocalliphora* blow flies either by acting as a physical barrier between larvae and nestlings or through chemical compounds associated with preen oil. However, empirical evidence for an antiparasitic effect of feathers remains equivocal. Using 1,100 naturally infested Tree Swallow (*Tachycineta bicolor)* nests collected from a nest-box study system monitored in southern Québec, Canada, between 2004 and 2018, we tested whether nest mass and feather abundance influenced hematophagous ectoparasites *(Protocalliphora* spp.) loads. We further assessed whether feather abundance directly affected body size using measurements of 1,513 *P. sialia* puparia, and emergence success using data from 961 *Protocalliphora* spp. puparia.

Nest mass was positively associated with feather abundance, and both variables showed very weak positive relationships with parasite abundance. Feather abundance was unrelated to puparium size nor parasite emergence success.

Our results provide no support for an anti-parasitic effect of feather lining within Tree Swallow nests. Using one of the largest datasets assembled to date on feather-lined nests and avian blow flies, we found no evidence that feather abundance impairs *Protocalliphora* density, development or emergence success. However, our findings do not preclude potential effects of feathers on other ectoparasites or microorganisms with different life histories. Future studies simultaneously monitoring multiple parasite taxa (such as fleas, mites, or bacteria) may therefore be needed to fully evaluate the ecological consequences of feather lining within avian nests.

## INTRODUCTION

Bird nests are among the most elaborate structures built by vertebrates and reflect multiple selective pressures acting on reproductive success. Nest have multiple functions including thermoregulation, structural stability, sexual selection, egg retention and predator avoidance (Hansell 1984, 2000, Garcia-Navas et al 2015). However, nests are more than breeding structures; they constitute dynamic microhabitats that can support diverse communities of organisms (Carvallo et al. 2020, Jaworski et al. 2022). Numerous arthropods, microorganisms and parasites exploit avian nests as shelter, food resources or breeding sites, creating complex networks of commensal, mutualistic and antagonistic interactions (Woodroffe 1953, Maziarz et al. 2022). Notably, nest-dwelling ectoparasites can influence nestling development, survival and reproductive success, although their effects often vary among host species and environmental contexts (Grab et al. 2019, Roby et al. 1992, Coroller-Chouraki et al., 2026b).

Birds can use a variety of defenses against ectoparasites. These include direct behavioral responses such as preening, parasite avoidance, and compensatory provisioning, whereby parents increase feeding rates to offset the energetic costs inflicted on parasitized nestlings (Christe et al. 1996, Simon et al. 2004, Bush & Clayton 2018). Many species also perform nest sanitation by actively removing waste material (Glądalski et al. 2025) or alter nest composition. For instance, birds may add aromatic green vegetation (Tomás et al 2012, Scott-Baumann and Morgan 2015, Glądalski et al. 2025) which possesses insecticidal or antimicrobial properties (Shutler and Campbell 2007, Clayton et al. 2010, Bush & Clayton 2018). Finally, the incorporation of human-derived materials (such as cigarette butts) or substrates like mammalian fur treated with veterinary parasiticides can successfully reduce ectoparasite loads, though often at the expense of higher toxicity and mortality for offspring (Suárez-Rodríguez et al. 2013, 2017, Tassin-de-Montaigu et al. 2026).

Among cavity-nesting passerines, Tree Swallows (*Tachycineta bicolor*) are known for incorporating large quantities of moulted feathers into their nests (Winkler 1993). This feather lining is widely considered adaptative: feathers improve nest insulation and can enhance reproductive performance (Møller 1984, Lombardo et al. 1995, but see Mainwaring et al 2016). Feathers are also thought to influence parasite-host interactions through physical barrier effects or antimicrobial properties mediated by uropygial oil coatings (Peralta-Sanchez et al. 2010, Mainwaring et al. 2014, Ruiz-Rodríguez et al. 2015). However, empirical support for these possible anti-parasitic benefits remains scarce (Moreno-Rueda 2017). In fact, feather incorporation may carry overlooked maladaptative costs depending on the parasite’s life history. Feathers coated with uropygial secretions may release complex volatile organic compounds into the nest environment, which can act as chemostimulants for calliphorid flies (Jeanbourquin & Guerin 2007). In line with this olfactory attraction, observational and experimental work has shown higher abundances of hematophagous *Protocalliphora* bird blowflies pupae in feather-rich nests (Dawson et al., 2011, Williams 2017). This challenges the anti-parasitic hypothesis, highlighting that current empirical evidence remains highly fragmented and contradictory. Yet, while feathers appear to attract ectoparasites, they might subsequently stunt the parasites’ growth and development (Williams 2017). This suggests that nest feathers may simultaneously enhance ectoparasite attraction while constraining their development (Winkler 1993; Williams 2017), leaving the net ecological outcome of this behavior unresolved due to a lack of large-scale studies.

To resolve these conflicting hypotheses, we used 1,100 Tree Swallow infested nests from a 10,200-km^2^ nest box study system monitored between 2004 and 2018 in Québec, Canada, and tested whether nest mass and feather abundance influenced *Protocalliphora* spp. bird blow flies load. Using data from individual *Protocalliphora* analysis, we then investigated whether feather abundance influenced *P. sialia* Shannon & Dobroscky puparia size, *Protocalliphora* spp. abundance and emergence success.

## Materials and methods

### Study system

We used data collected between 2004 and 2018 from a long-term nest-box study system established in southern Québec, Canada, to monitor Tree Swallows breeding in agricultural landscapes (Ghilain and Bélisle 2008). The study system comprises 400 nest boxes distributed among 40 farms spanning a 10,200-km^2^ gradient of agricultural intensification (10 nest boxes per farm).

### Agricultural, meteorological and nidification contexts

To isolate the effects of nest characteristics on *Protocalliphora* abundance, puparial size, and emergence success, we statistically controlled for ecological sources of variation already reported in Coroller-Chouraki et al., 2026b,c,d.

Landscape contexts were based on land-cover digitization within a 500m radius around nestboxes (QGIS), we performed a robust compositional PCA (rPCA) on yearly relative habitat areas (Forest, Corn/Soybean, Other Cereals, Pasture/Forage). Axis component 1 (Comp.1) reflects an intensification gradient (positive: maize/soybean; negative: forage/trees) and Axis component 2 (Comp.2) contrasts forest vs. forage cover. For 2004–2005, scores were extrapolated using 2006–2008 averages due to high landscape stability. Landscape contexts were approximated with the interaction between Comp.1 and Comp.2.

Meteorological contexts were approximated with the Mean hourly temperature (°C) and total precipitation (mm) extracted via the *NASAPOWER* R package using farm coordinates. For each nest, hourly data were averaged over the specific period spanning from the first hatching date to 7 days post-fledging/death to match *Protocalliphora* development and pupation. Nestling hatching date was estimated through the visit of each individual nest box every two days. Calendar date (Day of Year) of the first hatching event for each individual clutch, served as a proxy for seasonal phenology and the timing of initial ectoparasite access to nestling host blood. Nestling host availability served as a proxy for the total host biomass and duration of blood resource accessible to ectoparasites. It was quantified as the cumulative sum of ‘nestling-days’ per nest and calculated by summing the daily count of living nestlings present in the nest box each day from the first hatching event until fledging or total brood mortality.

### Nest sampling and *Protocalliphora* parasites

After each breeding season, we collected Tree Swallow nests in plastic bags and stored them in a cold room until processing. Nests originating from multiple breeding attempts within a season were excluded. Each nest was weighed to the nearest 0.01 g (P-2002, Denver instrument, Bohemia, NY, USA) and then dissected under a ventilated hood to recover and count all feathers as well as all *Protocalliphora* puparia which were subsequently preserved in 70% ethanol.

Species-level identification of *Protocalliphora* puparia was completed for a sub-sample of 216 nests (see Coroller-Chouraki et al. 2026a). Those puparia were examined under a stereomicroscope and identified using morphological keys (Sabrosky et al. 1989, Whitworth 2019). Although three *Protocalliphora* species were found in our samples (*P. sialia, P. bennetti* Whitworth and *P. metallica* Townsend), *P. sialia* largely dominated infestations (> 80% of identified puparia; Coroller-Chouraki et al. 2026a). Because species-level identification of all puparia was not feasible and because all three species share a similar parasitic larval ecology as hematophagous nest ectoparasites of altricial birds (Bennett & Whitworth 1991), all *Protocalliphora* puparia were pooled for analyses concerning abundance and emergence.

To assess parasite size, all puparia were photographed under a calibrated stereomicroscope. Morphometric measurements were extracted using an automated Fiji/ImageJ workflow following Coroller-Chouraki et al. (2026a). Because *P. sialia* represented most specimens in our subsample and exhibited consistent occurrence throughout the study period, analyses of parasite size were restricted to this species. For each image, a threshold-based segmentation isolated individual puparia, and the “Fit Ellipse” function was applied to each object. This method modelled each *P. sialia* puparium as an ellipsoid and calculated its theoretical surface area (mm^2^), which we used as a proxy for size. Measuring this projected area rather than length or width minimizes bias due to the missing operculum in emerged puparia, while providing a reproducible 2D metric that reflects both elongation and width variation.

We estimated *Protocalliphora* spp. emergence success under a stereomicroscope. In our study system, parasitoid wasps of the genus *Nasonia* (Hymenoptera: Pteromalidae) constitute the primary source of *Protocalliphora* mortality, accounting on average for 68.8% of bird blowflies deaths and reaching up to 90.5% in some years (Coroller-Chouraki et al. 2026a). For each *Protocalliphora* individual included in the subsample, evidence of *Nasonia* parasitism was recorded, including parasitoid emergence holes or the presence of unemerged *Nasonia* individuals within dissected puparia. Failed *Protocalliphora* emergence unrelated to parasitoid attack was identified by the presence of emergence caps and pupal remains without evidence of *Nasonia* development. To isolate the potential effect of feather abundance on parasite emergence success, all *Protocalliphora* individuals whose mortality was directly attributable to *Nasonia* parasitism (n = 2,142 out of 3,103 individuals) were excluded, leaving out 961 uninfested *Protocalliphora* individuals for the analysis.

### Statistical analyses

First, we analyzed the statistical relationships between nest mass and feather abundance. Then, we tested whether nest mass and/or feather abundance predicted (i) *Protocalliphora* abundance, and (ii) *P. sialia* puparium size, controlling for host availability, breeding phenology, meteorological conditions and landscape context, following Cinelli et al. (2022) recommendations. We used *glmmTMB* (v.1.1.9; Brooks et al. 2017) or *mgcv* (v.1.8.42; Wood 2023) R packages and generated model predictions at the population-level by marginalizing over the random effect structure and setting non-focal explanatory variables at their means using the *marginaleffects* R package (v.0.18.0, Arel-Bundock, 2024). The effects of these statistical controls have been examined in detail elsewhere (Coroller-Chouraki et al. 2026 b,d). Farm identity and year were included as nested random effects to account for spatial and temporal non-independence (Tab. 1). The conditional and marginal *R*^2^ (respectively R^2^c and R^2^m) of the GLMM were calculated following Nakagawa’s et al. (2017) with the *MuMIn* package (v.1.47.5, Barton 2023). Given that random effects are treated as smooth term, which complicates variance partitioning in GAMMs we only provide the adjusted *R*^2^ returned by the *mgcv* summary function (Wood, 2023). Seven nests from farms that were located particularly close to poultry industries resulted in an extremely high number of feathers (up to 500) being found in the lining (compared to an overall median of 26 feathers in nests infested by *Protocalliphora*). These seven nests caused outlier-driven results and were excluded from the analysis *a posteriori*.

**Table 1.**
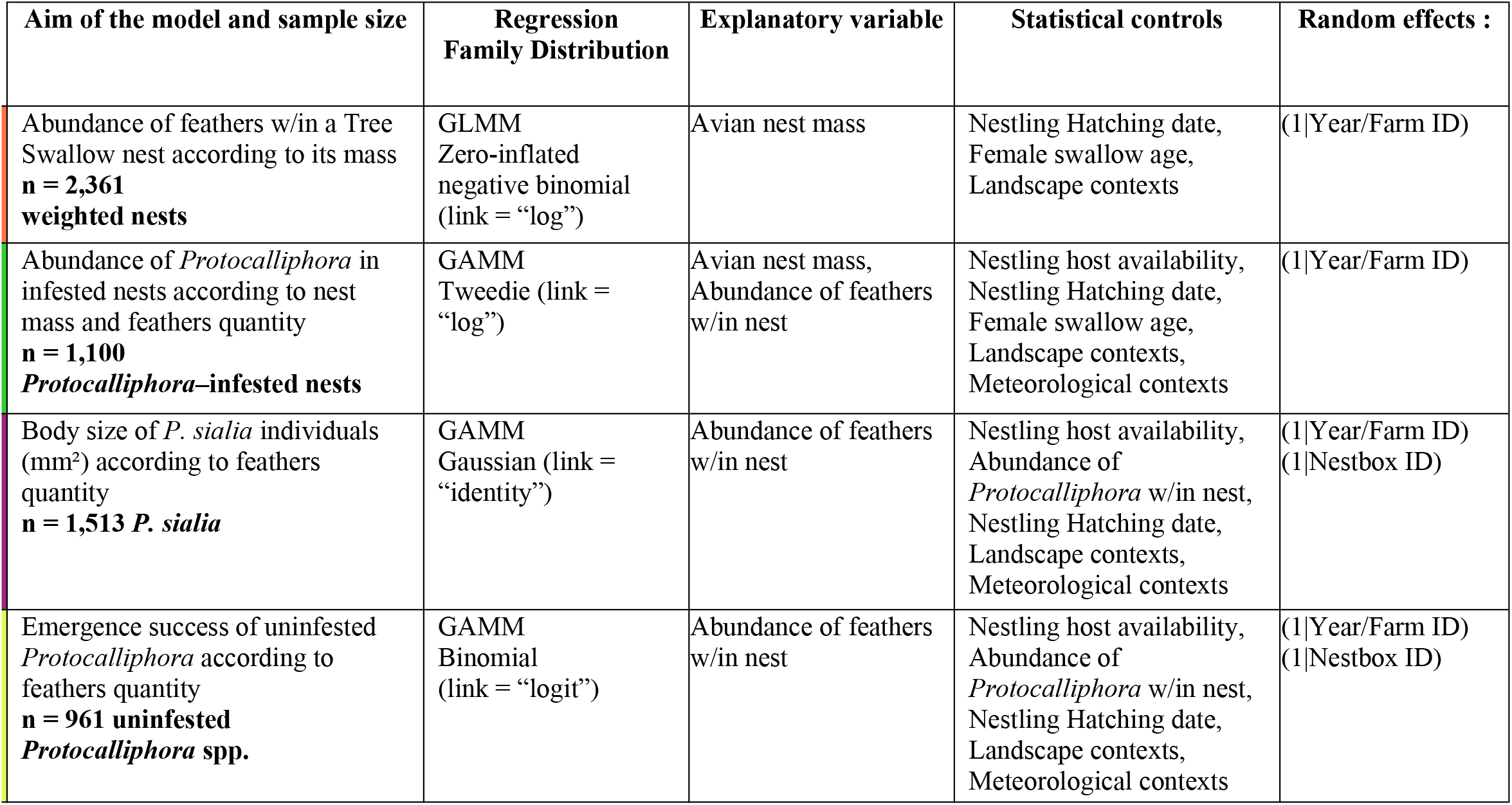
Models used to estimate the direct effects between Tree Swallow nest mass, feather quantity, *Protocalliphora* spp. density and emergence success, and *P. sialia* puparial size.

| Aim of the model and sample size | Regression Family Distribution | Explanatory variable | Statistical controls | Random effects : |
| --- | --- | --- | --- | --- |
| Abundance of feathers w/in a Tree Swallow nest according to its mass<br>**n = 2,361 weighted nests** | GLMM<br>Zero-inflated negative binomial (link = “log”) | Avian nest mass | Nestling Hatching date, Female swallow age, Landscape contexts | (1\|Year/Farm ID) |
| Abundance of *Protocalliphora* in infested nests according to nest mass and feathers quantity<br>**n = 1,100 Protocalliphora–infested nests** | GAMM<br>Tweedie (link = “log”) | Avian nest mass, Abundance of feathers w/in nest | Nestling host availability, Nestling Hatching date, Female swallow age, Landscape contexts, Meteorological contexts | (1\|Year/Farm ID) |
| Body size of *P. sialia* individuals (mm<sup>2</sup>) according to feathers quantity<br>**n = 1,513 *P. sialia*** | GAMM<br>Gaussian (link = “identity”) | Abundance of feathers w/in nest | Nestling host availability, Abundance of *Protocalliphora* w/in nest, Nestling Hatching date, Landscape contexts, Meteorological contexts | (1\|Year/Farm ID)<br>(1\|Nestbox ID) |
| Emergence success of uninfested *Protocalliphora* according to feathers quantity<br>**n = 961 uninfested Protocalliphora spp.** | GAMM<br>Binomial (link = “logit”) | Abundance of feathers w/in nest | Nestling host availability, Abundance of *Protocalliphora* w/in nest, Nestling Hatching date, Landscape contexts, Meteorological contexts | (1\|Year/Farm ID)<br>(1\|Nestbox ID) |

## RESULTS

Nest mass was positively associated with feather abundance, indicating that larger nests generally contained more feathers (Fig. 1a). However, the effect size was relatively small (β = 0.101 ± 0.020 SE, P < 0.001). Marginal (R^2^m) and conditional (R^2^c) R^2^ values suggested that a larger proportion of the explained variation in feather abundance was due to random effects (farm and year; R^2^m = 0.026–0.032, R^2^c = 0.182–0.220). Female age (second-year versus after-second-year) did not influence feather abundance within nests.

**Figure 1.**
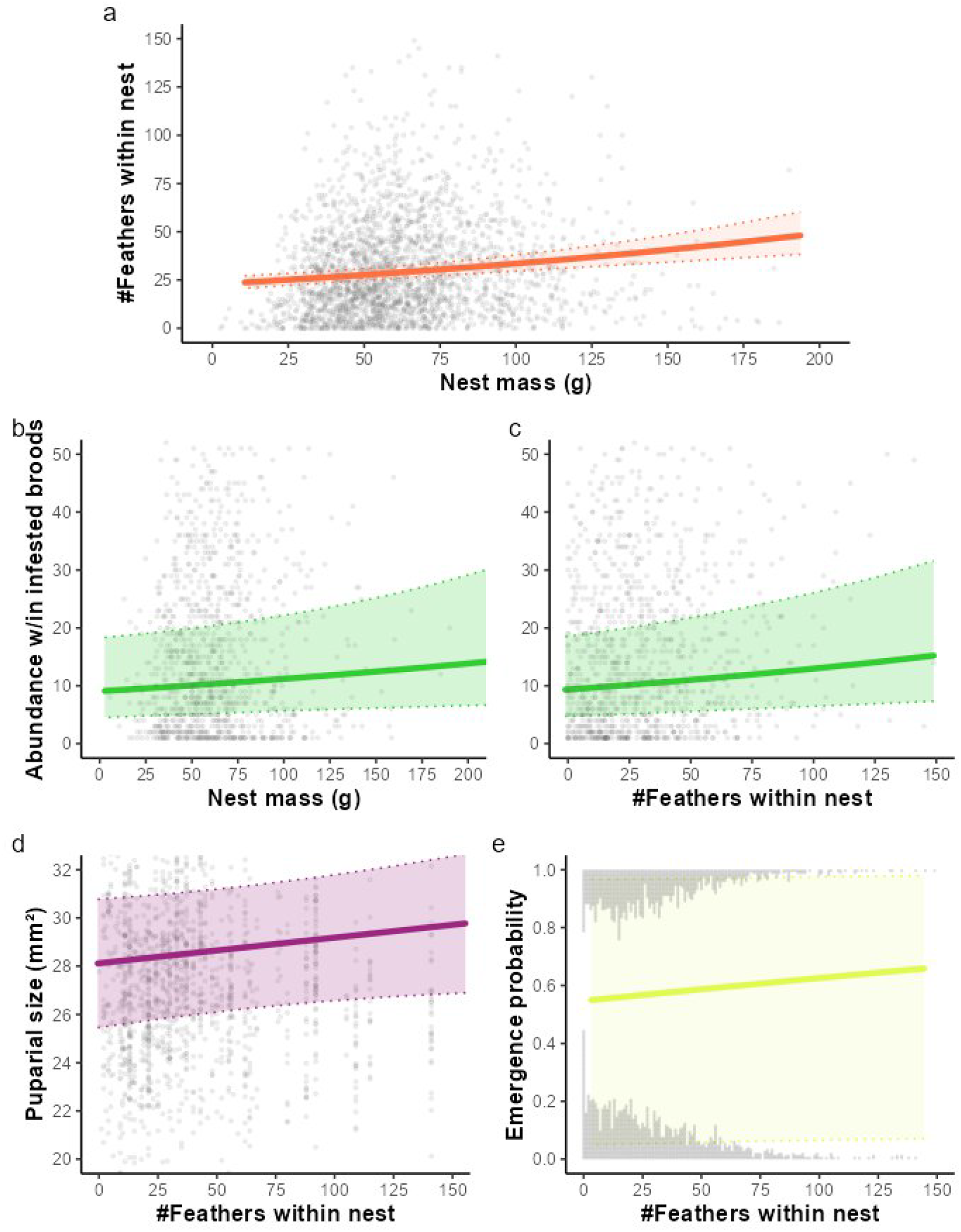
Variation in a) feathers abundance within a nest according to nest mass (g); b) and c) abundance of *Protocalliphora* spp. within infested broods according to nest mass and feathers abundance within the nest; d) *P. sialia* puparial size (mm^2^) according to feathers abundance within the nest, e) *Protocalliphora* spp. emergence probability according to feathers abundance within the nest. Grey dots/lines indicate raw data.

Among nests infested by *Protocalliphora* spp., parasite abundance increased slightly with both nest mass and feather abundance (Fig. 1b, c). Model (adjusted R^2^ = 0.256) showed the effect of nest mass to be weak and only marginally non-significant (β = 0.055 ± 0.031 SE, P = 0.079), whereas feather abundance had a weak significant positive effect on parasite abundance (β = 0.082 ± 0.030 SE, P = 0.006).

Feather abundance was not associated with the size of *P. sialia* puparia (Fig. 1d; β = 0.332 ± 0.219 SE, P = 0.131), despite the model explaining a substantial proportion of variation in puparium size (adjusted R^2^ = 0.365).

Similarly, feather abundance had no effect on the emergence probability of *Protocalliphora* puparia after excluding individuals parasitized by *Nasonia* spp. (Fig. 1e; β = 0.091 ± 0.131 SE, P = 0.484). The model explained 26.6% of the variation in emergence success, which was primarily associated with host availability and hatching phenology.

## DISCUSSION

The positive relationship between nest mass and the quantity of feathers incorporated within suggests individual behavioral plasticity in nest building effort, where higher-quality individuals construct heavier structures and collect more feathers (Deeming et al 2019; Järvinen & Brommer 2020). Conversely, individuals facing high parasite loads or poorer physiological condition often exhibit reduced construction investment, building lighter nests with fewer feathers (Tomás et al. 2006).

More importantly, our results provide no support for the hypothesis that feather incorporation in Tree Swallow nests acts as a structural or chemical defense against *Protocalliphora*. While previous experimental work reported either an increase in parasite loads (Dawson et al., 2011) or no change in arthropod composition (Stephenson et al. 2009), no change in Hen Fleas (*Ceratophyllus gallinae*) load within Blue Tit (*Cyanistes caeruleus*) nests, we found a negligible, near-zero relationship between bird blow fly abundance and feather quantity or Tree Swallow nest mass. It is unlikely that the olfactory cues attracting gravid females originate from these materials. Furthermore, feather abundance did not influence *P. sialia* puparial size or *Protocalliphora* spp. emergence success. This absence of developmental costs on larval traits within feather-rich nests challenges hypotheses suggesting that feathers, or the preen oil they carry, impair parasite fitness (e.g., Williams 2017). Given the unprecedented scale of our longitudinal study (n = 1,100 infested nests) compared to previous investigations (n = 62 in Dawson et al. 2011; n = 55 in Stephenson et al. 2009; n = 45 in Williams 2017), our results indicate that *Protocalliphora* load and fitness are not driven by nest architecture. Instead, abundance and performance appear mainly induced by other ecological drivers, such as density-dependent interactions among hosts, parasites, and parasitoids, alongside phenological, meteorological, and landscape conditions (Musgrave et al., 2019, Garrido-Bautista et al. 2020, Castaño-Vázquez et al. 2021, Mennerat et al. 2021, Merino et al. 2023, Coroller-Chouraki et al. 2026b,d).

The apparent lack of effect observed in our study should not be generalized to the entire nest ecosystem as nests shelter complex communities of arthropods and microorganisms that may interact with nest lining material in ways yet to be uncovered (Mainwaring et al. 2014, Ruiz-Rodríguez et al. 2015, Moreno-Rueda 2017). Feather lining could still modulate other community components through physical, chemical, or microbial pathways (but see Stephenson et al. 2009).

For instance, feathers alter the microclimate within the nest box which are known to affect parasite dynamics and host-parasite balances (Mennerat et al. 2021, Merino et al. 2023). Ultimately, if an anti-parasitic role is discounted, it suggests that feather incorporation is maintained by a trade-off between social signaling benefits (Tomás et al, 2012; Garcia-Navas et al 2015) and the energetic costs associated, notwithstanding their potential function in nest insulation. Adopting a broader community-level approach in future studies remains essential to fully untangle the multi-trophic consequences of nest-building behaviors.

## REFERENCES

Arel-Bundock, V., N. Greifer, and A. Heiss. 2024. How to interpret statistical models using marginaleffects for R and Python. Journal of Statistical Software 111:1–32.

Bartoń, K. 2026. MuMIn: Multi-Model Inference. R package version 1.48.19. https://CRAN.R-project.org/package=MuMIn

Bennett, G. F., and T. L. Whitworth. 1991. Studies on the life history of some species of Protocalliphora (Diptera: Calliphoridae). Canadian Journal of Zoology 69:2048–2058.

Brooks, M. E., K. Kristensen, K. J. van Benthem, A. Magnusson, C. W. Berg, A. Nielsen, H. J. Skaug, M. Maechler, and B. M. Bolker. 2017. glmmTMB balances speed and flexibility among packages for zero-inflated generalized linear mixed modeling. The R Journal 9:378–400.

Bush, S. E., and D. H. Clayton. 2018. Anti-parasite behaviour of birds. Philosophical Transactions of the Royal Society B: Biological Sciences 373:20170196.

Carvallo, G. O., M. López-Aliste, M. Lizama, N. Zamora, and G. Muschett. 2020. Assessing climatic and intrinsic factors that drive arthropod diversity in bird nests. Gayana 84:16–27.

Castaño-Vázquez, F., Y. R. Schumm, A. Bentele, P. Quillfeldt, and S. Merino. 2021. Experimental manipulation of cavity temperature produces differential effects on parasite abundances in blue tit nests at two different latitudes. International Journal for Parasitology: Parasites and Wildlife 14:287–297.

Christe, P., H. Richner, and A. Oppliger. 1996. Begging, food provisioning and nestling competition in great tit broods infested with ectoparasites. Behavioral Ecology 7:127–131.

Cinelli, C., A. Forney, and J. Pearl. 2024. A crash course in good and bad controls. Sociological Methods & Research 53:1071–1104.

Clayton, D. H., J. H. Koop, C. W. Harbison, B. R. Moyer, and S. E. Bush. 2010. How birds combat ectoparasites. Open Access Publications, University of Nebraska-Lincoln.

Coroller-Chouraki, S., J. Savage, F. Pelletier, D. Garant, and M. Bélisle. 2026a. Parasitism by, species composition, morphometry, and parasitoidism of Protocalliphora bird blowflies (Diptera: Calliphoridae) in Quebec’s farmlands: A 16-year dataset. The Canadian Entomologist 158:e11.

Coroller-Chouraki, S., A. Bush-Beaupré, J. Savage, and M. Bélisle. 2026b. Enemy mitigation in farmlands: Intensive agriculture impacts avian nesting and subsequent ectoparasite infestation. bioRxiv 2026.02.

Coroller-Chouraki, S., J. Savage, and M. Bélisle. 2026c. Density-dependence mechanisms and agricultural intensification affect host-ectoparasite-hyperparasite interactions in a bird-insect non-targeted tri-trophic system. bioRxiv 2026.10.

Coroller-Chouraki, S., J. Savage, and M. Bélisle. 2026d. Selective survival of avian ectoparasites in agrointensive habitats and subsequent effects on host selection by parasitoids. bioRxiv 2026.10.

Dawson, R. D., C. C. O’Brien, and K. L. Malyon. 2011. The price of insulation: costs and benefits of feather delivery to nests for male tree swallows Tachycineta bicolor. Journal of Avian Biology 42:394–402.

Deeming, D. C., F. E. Morton, and K. L. Laverack. 2019. Nestbox size affects mass and proportions of materials used in blue tit Cyanistes caeruleus nests. Bird Study 66:130–135.

García-Navas, V., F. Valera, and M. Griggio. 2015. Nest decorations: an ‘extended’ female badge of status? Animal Behaviour 99:95–107.

Garrido-Bautista, J., G. Moreno-Rueda, A. Baz, D. Canal, C. Camacho, B. Cifrián, J. L. Nieves-Aldrey, M. Carles-Tolrá, and J. Potti. 2020. Variation in parasitoidism of Protocalliphora azurea (Diptera: Calliphoridae) by Nasonia vitripennis (Hymenoptera: Pteromalidae) in Spain. Parasitology Research 119:559–566.

Ghilain, A., and M. Bélisle. 2008. Breeding success of tree swallows along a gradient of agricultural intensification. Ecological Applications 18:1140–1154.

Glądalski, M., A. C. Norte, M. Bartos, I. Demeško, A. Kaliński, M. Markowski, J. Skwarska, J. Wawrzyniak, P. Zieliński, and J. Bańbura. 2025. Effects of experimental nest treatment with herbs on ectoparasites and body condition of nestlings. Behavioral Ecology 36:arae103.

Grab, K. M., B. J. Hiller, J. H. Hurlbert, M. E. Ingram, A. B. Parker, D. Y. Pokutnaya, and S. A. Knutie. 2019. Host tolerance and resistance to parasitic nest flies differs between two wild bird species. Ecology and Evolution 9:12144–12155.

Hansell, M. H. 1984. Animal Architecture and Building Behaviour. Longman, lLondon.

Hansell, M. H. 2000. Bird Nests and Construction Behaviour. Cambridge University Press, Cambridge.

Järvinen, P., and J. E. Brommer. 2020. Lining the nest with more feathers increases offspring recruitment probability: Selection on an extended phenotype in the blue tit. Ecology and Evolution 10:13327–13333.

Jaworski, T., J. Gryz, D. Krauze-Gryz, R. Plewa, C. Bystrowski, R. Dobosz, and J. Horák. 2022. My home is your home: Nest boxes for birds and mammals provide habitats for diverse insect communities. Insect Conservation and Diversity 15:461–469.

Jeanbourquin, P., and P. M. Guerin. 2007. Sensory and behavioural responses of the stable fly Stomoxys calcitrans to rumen volatiles. Medical and Veterinary Entomology 21:217–224.

Lombardo, M. P., R. M. Bosman, C. A. Faro, S. G. Houtteman, and T. S. Kluisza. 1995. Effect of feathers as nest insulation on incubation behavior and reproductive performance of Tree Swallows (Tachycineta bicolor). The Auk 112:973–981.

Mainwaring, M. C., I. R. Hartley, M. M. Lambrechts, and D. C. Deeming. 2014. The design and function of birds’ nests. Ecology and Evolution 4:3909–3928.

Mainwaring, M. C., A. Wolfenden, J. E. Read, J. M. Robson, C. J. Tomlinson, and I. R. Hartley. 2016. Feathering the nest: the effects of feather supplementation to blue tit nests. Avian Biology Research 9:89–95.

Maziarz, M., R. K. Broughton, P. Chylarecki, and G. Hebda. 2022. Weather impacts on interactions between nesting birds, nest-dwelling ectoparasites and ants. Scientific Reports 12:17845.

Mennerat, A., A. Charmantier, P. Perret, S. Hurtrez-Boussès, and M. M. Lambrechts. 2021. Parasite intensity is driven by temperature in a wild bird. Peer Community Journal 1:e60.

Merino, M., M. García-del-Río, F. Castaño-Vázquez, and S. Merino. 2023. A long-term study on the influence of climate on two common nest-dwelling ectoparasites, the blowfly pupae Protocalliphora azurea and the mite Dermanyssus gallinoides in nests of the Eurasian Blue tit (Cyanistes caeruleus: Aves, Passeriformes, Paridae). Parasitology Research 122:1500–1512.

Møller, A. P. 1984. On the use of feathers in birds’ nests: predictions and tests. Ornis Scandinavica 15:38–42.

Moreno-Rueda, G. 2017. Preen oil and bird fitness: a critical review of the evidence. Biological Reviews 92:2131–2143.

Musgrave, K. A., R. D. Dawson, and C. C. O’Brien. 2019. Long-term variation in environmental conditions influences host–parasite fitness. Ecology and Evolution 9:5321–5330.

Nakagawa, S., P. C. D. Johnson, and H. Schielzeth. 2017. The coefficient of determination R2 and intra-class correlation coefficient from generalized linear mixed-effects models revisited and expanded. Journal of The Royal Society Interface 14:20170213.

Peralta-Sánchez, J. M., A. P. Møller, A. M. Martín-Platero, and J. J. Soler. 2010. Number and colour composition of nest lining feathers predict eggshell bacterial community in barn swallow nests: an experimental study. Functional Ecology 24:426–433.

Roby, D. D., K. L. Brink, and K. Wittmann. 1992. Effects of bird blowfly parasitism on eastern bluebird and tree swallow nestlings. The Wilson Bulletin 104:630–643.

Ruiz-Rodríguez, M., G. Tomás, D. Martín-Gálvez, C. Ruiz-Castellano, and J. J. Soler. 2015. Bacteria and the evolution of honest signals: The case of ornamental throat feathers in spotless starlings. Functional Ecology 29:701–709.

Sabrosky, C. W., G. F. Bennett, and T. L. Whitworth. 1989. Bird blow flies (Protocalliphora) in North America (Diptera: Calliphoridae), with notes on Palearctic species. Smithsonian Institution Press, Washington, D.C.

Scott-Baumann, J. F., and E. R. Morgan. 2015. A review of the nest protection hypothesis: does inclusion of fresh green plant material in birds’ nests reduce parasite infestation? Parasitology 142:1016–1023.

Shutler, D., and A. A. Campbell. 2007. Experimental addition of greenery reduces flea loads in nests of a non-greenery using species, the tree swallow Tachycineta bicolor. Journal of Avian Biology 38:7–12.

Simon, A., D. Thomas, J. Blondel, P. Perret, and M. M. Lambrechts. 2004. Physiological ecology of Mediterranean Blue Tits (Parus caeruleus L.): effects of ectoparasites (Protocalliphora spp.) and food abundance on metabolic capacity of nestlings. Physiological and Biochemical Zoology 77:492–501.

Stephenson, S., S. Hannon, and H. Proctor. 2009. The function of feathers in Tree Swallow nests: insulation or ectoparasite barrier? The Condor 111:479–487.

Suárez-Rodríguez, M., I. López-Rull, and C. Macías Garcia. 2013. Incorporation of cigarette butts into nests reduces nest ectoparasite load in urban birds: new ingredients for an old recipe? Biology Letters 9:20120931.

Suárez-Rodríguez, M., R. D. Montero-Montoya, and C. Macías Garcia. 2017. Anthropogenic nest materials may increase breeding costs for urban birds. Frontiers in Ecology and Evolution 5:4.

Tassin-de-Montaigu, C., G. Glauser, S. Guinchard, and D. Goulson. 2026. Transfer of veterinary parasiticides from the fur lining bird’s nest to eggs and chicks. Environmental Science and Pollution Research 33:7070–7081.

Tomás, G., S. Merino, J. Moreno, J. J. Sanz, J. Morales, and S. García-Fraile. 2006. Nest weight and female health in the blue tit (Cyanistes caeruleus). The Auk 123:1013–1021.

Tomás, G., S. Merino, J. Martínez-de la Puente, J. Moreno, J. Morales, E. Lobato, S. Rivero-de Aguilar, and S. Del Cerro. 2012. Interacting effects of aromatic plants and female age on nest-dwelling ectoparasites and blood-sucking flies in avian nests. Behavioural Processes 90:246–253.

Whitworth, T. L. 2019. Keys to the genera and species of blow flies (Diptera: Calliphoridae) of America, North of Mexico. In: Forensic Entomology. CRC Press, Boca Raton, FL, pp. 413–443.

Williams, D. 2017. Parasite-host interactions between Protocalliphora Spp. (Diptera: Calliphoridae) and tree swallows Tachycineta bicolor. Master’s thesis, University of Northern British Columbia, Prince George, Canada.

Winkler, D. W. 1993. Use and importance of feathers as nest lining in Tree Swallows (Tachycineta bicolor). The Auk 110:29–36.

Wood, S. N. 2023. mgcv: Mixed GAM Computation Vehicle with Automatic Smoothness Estimation. R package version 1. 9–0. https://CRAN.R-project.org/package=mgcv

Woodroffe, G. E. 1953. An ecological study of the insects and mites in the nests of certain birds in Britain. Bulletin of Entomological Research 44:739–772.

